# Systematic Optimization of Whole Plant Carbon Nitrogen Interaction (WACNI) to Support Crop Design for Greater Yield

**DOI:** 10.1101/286112

**Authors:** Tian-Gen Chang, Xin-Guang Zhu

**Affiliations:** CAS-MPG Partner Institute for Computational Biology, Shanghai Institutes for Biological Sciences, Chinese Academy of Sciences, Shanghai 200031, China; University of Chinese Academy of Sciences, 19A Yuquan Road, Beijing 100049, China; Key Laboratory for Plant Molecular Genetics, Center of Excellence for Molecular Plant Sciences, Chinese Academy of Sciences, Shanghai 200031, China

**Keywords:** crop yields, grain filling, molecular breeding, source sink interaction, systems model

## Abstract

On the face of the rapid advances in genome editing technology and greatly expanded knowledge on plant genome and genes, there is a strong demand to develop an effective tool to guide designing crops for higher yields. Here we developed a highly mechanistic model of Whole plAnt Carbon Nitrogen Interaction (WACNI), which predicts crop yield based on major metabolic and biophysical processes in source, sink and transport tissues. WACNI accurately predicted the yield responses of so far reported source, sink and transport related genetic manipulations on rice grain yields. Systematic sensitivity analysis with WACNI was used to classify the source, sink and transport related molecular processes into four categories, i.e. universal yield enhancers, universal yield inhibitors, conditional yield enhancers and weak yield regulators. Simulations using WACNI further show that even without a major change in leaf photosynthetic properties, 54.6% to 73% grain yield increase can be potentially achieved by optimizing these molecular processes during the rice grain filling period while simply combining all the ‘superior’ molecular modules together cannot achieve the optimal yield level. A common macroscopic feature in all these designed high-yield lines is that they all show ‘a sustained and steady growth of grain sink’, which might be used as a generic selection criteria in high-yield rice breeding. Overall, WACNI can serve as a tool to facilitate plant source sink interaction research and guide future crops breeding by design.

**One sentence summary:** A mechanistic model of source, sink flow model is developed and used to demonstrate that optimization of the whole plant carbon nitrogen metabolism can dramatically increase crop yield potential.

## INTRODUCTION

With great advance achieved in functional genomics studies in major crops, phenomics and genome editing technologies (Araus and Cairns, 2014; Hartung and Schiemann, 2014; Sander and Joung 2014), rational crop design followed by genome editing are playing an increasingly important role in crop improvements for higher yield, better quality and enhanced resistance to environmental stresses (Wang et al., 2014a; Mickelbart et al., 2015; Qian et al., 2016). The challenge now is to develop highly mechanistic crop growth and developmental models, which can accurately predict the growth and development of crops, and correspondingly the crop yields and quality after one or a number of genes are manipulated in a particular cultivar (Hu and Xiong, 2014; Long et al., 2015; Xiao et al., 2017). Such models need to capture the non-linear properties of crop yield, quality and environmental responses, not from empirical correlation based prediction but rather from emergent behavior out of the interaction of the basic processes in the complex crop source, sink and transport systems (Chang et al., 2017).

Ever since the introduction of the first crop growth model by de Wit (1965), a large number of *in silico* models have been developed (Thornley, 1972; Minchin et al., 1993; Allen et al., 2005; Yin and van Laar, 2005), these models lack sufficient molecular details and hence cannot play the long-desired role of guiding crop design and breeding. This is not surprising since the foundation of most contemporary crop models were laid in a time when the understanding of molecular processes underlying crop growth and development was limited, and also the computational capacity was constrained which necessitated development of highly simplified models. Zhu et al (2015) provided strong evidence suggesting that it is feasible now to develop such highly mechanistic models of crop growth and development. Briefly, much progress has been made in developing many component modules for such a complete model, such as highly mechanistic models of leaf photosynthesis (Zhu et al., 2013), flowering time determination (Valentim et al., 2015), root growth and development (Lynch et al., 1997), and 3D canopy light interception (Song et al., 2013). Missing in the current repertoire of models is a highly mechanistic model describing metabolites transport and partitioning between source and sink organs and its consequence on plant growth and development. This problem has been dealt with in earlier models by various empirical approaches, ranging from simply using partitioning table or empirical equations (Aggarwal et al., 1994; Rasse and Tocquin, 2006), to assuming certain balanced partitioning of assimilates into different organs (Yin and van Laar, 2005). As a consequence of the empirical nature of these models, they may produce unreliable predictions under conditions where they have not been parameterized for. This is clearly reflected in a recent model comparison study where predicted impacts of elevated CO_2_ and temperature on rice yield with 13 models range from all the way from positive to negative (Li et al., 2015a).

Here we report the development, validation and application of a mechanistic model of Whole-plAnt Carbon and Nitrogen Interaction (WACNI), which can accurately predict the commonly phenomena related to source, sink and transport tissues. The WACNI model achieved this high accuracy by describing at a molecular scale the major biochemical and biophysical processes in source, sink and transport tissues during the grain filling period, when extensive non-linear interaction between these tissues occurs and the final crop yields are largely determined. WACNI was developed as a generic model, i.e. it can be parameterized to model different crops; here to take advantage of the large volumes of genetic and physiological studies available for rice, we first parameterized WACNI for rice. This paper describes the structure of the WACNI model, its parameterization, and its application in guiding optimization of source sink interaction to increased crop yields.

## RESULTS

### Features and performance of WACNI: a highly mechanistic model of source sink interaction

WACNI comprises five sub-models, i.e. root, leaf, grain, culm storage pool (CSP) and xylem-phloem transport system (Fig. 1). Fourteen major biochemical and biophysical processes involved in different source/sink/transport organs are described in WACNI. These processes included assimilation, transport, utilization and interaction of six primary metabolites, i.e. triose phosphates (TP), sucrose (Suc), starch, inorganic N (I-N, including NH4^+^ and NO_3_^-^), free form organic N (O-N, including amino acids and amides) and protein (Fig. 1).

**Figure 1.**
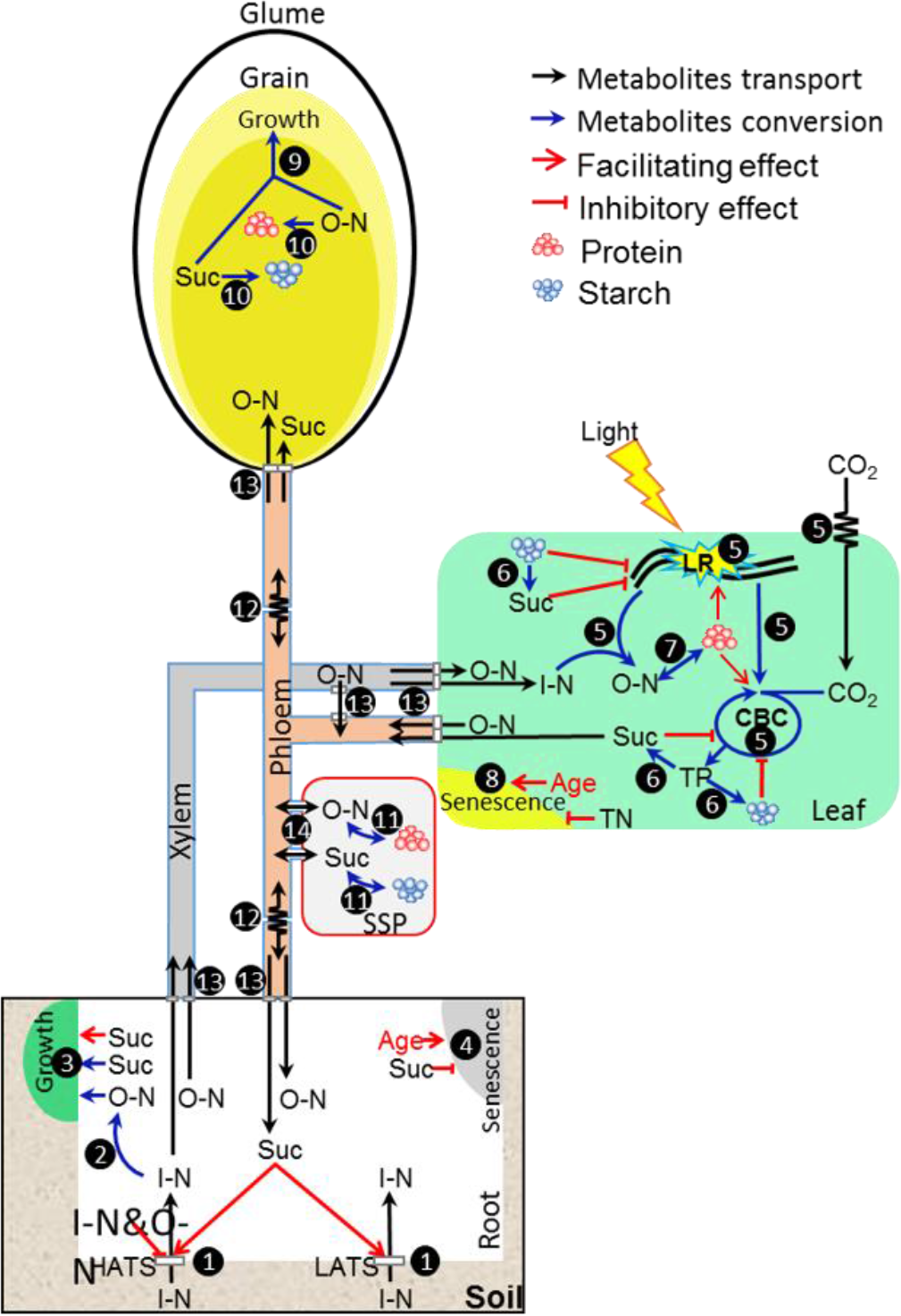
Diagram showing the biochemical/biophysical processes described in the WACNI model. These processes are classified into 14 groups which are shown in closed circles, they are: 1, root I-N uptake; 2, root I-N assimilation; 3, root growth; 4, root senescence; 5, leaf photosynthetic CO_2_ and I-N assimilation; 6, leaf triose phosphate, sucrose and starch interconversion; 7, leaf organic N and protein interconversion; 8, leaf senescence; 9, grain volume growth (cell division); 10, grain starch and protein accumulation; 11, culm storage pool sucrose and starch, O-N and protein interconversion; 12, osmotic pressure driven long distance phloem transport of sucrose and O-N; 13, transporter-dependent short distance phloem transport of sucrose and O-N (xylem and phloem loading and unloading, xylem-to-phloem transfer); 14, symplastic diffusion between phloem and culm storage pool. Abbreviations: Suc, sucrose; TP, triose phosphates; I-N, inorganic nitrogen; O-N, free-form organic nitrogen; HATS, high-affinity nitrogen transport system; LATS, low-affinity nitrogen transport system; CSP: culm storage pool; LR: light reaction; CBC, Calvin-Benson cycle.

After WACNI was parameterized for rice (Table 1; see a complete parameter list and description in Supplemental Table S1), it not only reached a steady state, but also realistically simulated many of the commonly observed dynamic changes during the rice grain filling period in root (Fig. 2a-d), leaf (Fig. 2e-h), grain (Fig. 2i-l) and culm (Fig. 2m-p). The predictions include changes of C and N assimilation rate in source organs (Fig. 2c, f), substrates concentration or content in different organs (Fig. 2d, g, h, l, m-p), organ growth and senescence (Fig. 2a, e, i, k) and organ respiration rates (Fig. 2b, j). The predicted trends are highly consistent with experimental data from literature (gray dots in Fig. 2e, f, h, i-l, p). These results suggest that despite the high complexity of potential signaling and regulatory mechanisms related to source sink interaction, the current known biochemical reactions, biophysical processes and their interactions incorporated in the model can already largely underlie the commonly observed rice growth and developmental patterns during the grain filling period.

**Figure 2.**
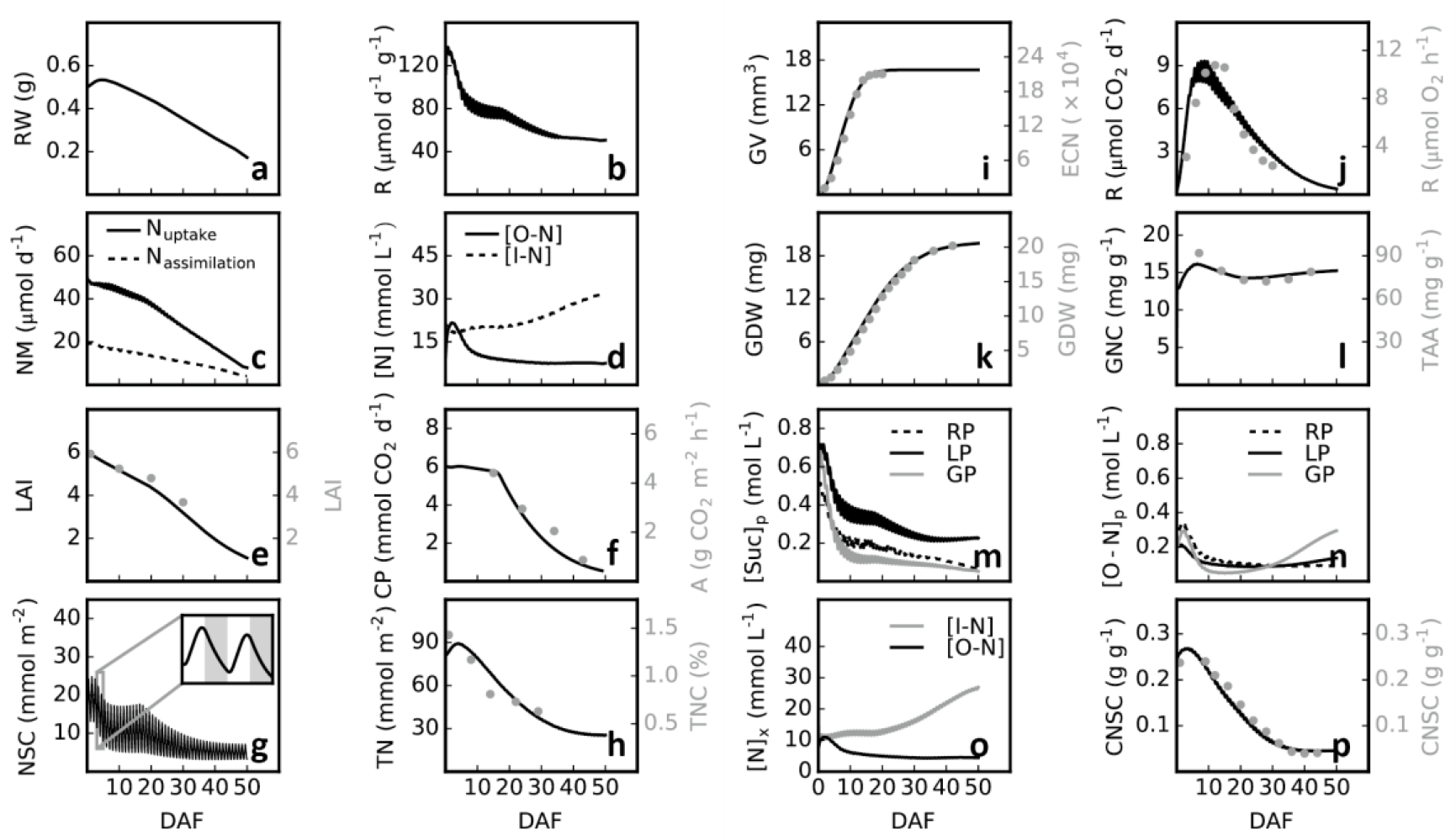
WACNI predictions of major metabolic and physiological changes in different organs, i.e. root, leaf, grain and culm, during the grain filling period after it’s parameterized for rice. Root (**a-d**): **a**, dry weight; **b**, respiration rate; **c**, I-N uptake and assimilation rate; **d**, O-N and I-N concentrations. Leaf (**e-h**): **e**, modeled and measured leaf area index (LAI) (Haque et al., 2015); **f**, modeled and measured canopy photosynthesis (Zhao et al., 2001); **g**, diurnal change of leaf non-structural carbohydrates concentration; **h**, modeled and measured leaf total nitrogen content (Dingkuhn et al., 1991). Grain (**i-l**): **i**, modeled grain volume (GV) increase and measured endosperm cell number (ECN) increase after flowering (Yang et al., 2006); **j**, modeled and *in vitro* measured grain respiration rate (Chen et al., 2006); **k**, modeled and measured grains dry weight (GDW) increase (Yang et al., 2006); **l**, modeled grain total N concentration (GNC) and measured total amino acids (TAA) concentration (Zhao et al., 2015). Culm (**m-p**): **m**, sucrose concentration ([Suc]_p_) in phloem compartment of root (RP), leaf (LP) and grain (GP); **n**, O-N concentration ([O-N]_p_) in phloem compartment of root (RP), leaf (LP) and grain (GP); **o**, I-N and O-N concentration in xylem; **p**, modeled and measured culm non-structural carbohydrates content (CNSC) (Yang et al., 2001). DAF: day after flowering.

**Table 1.**
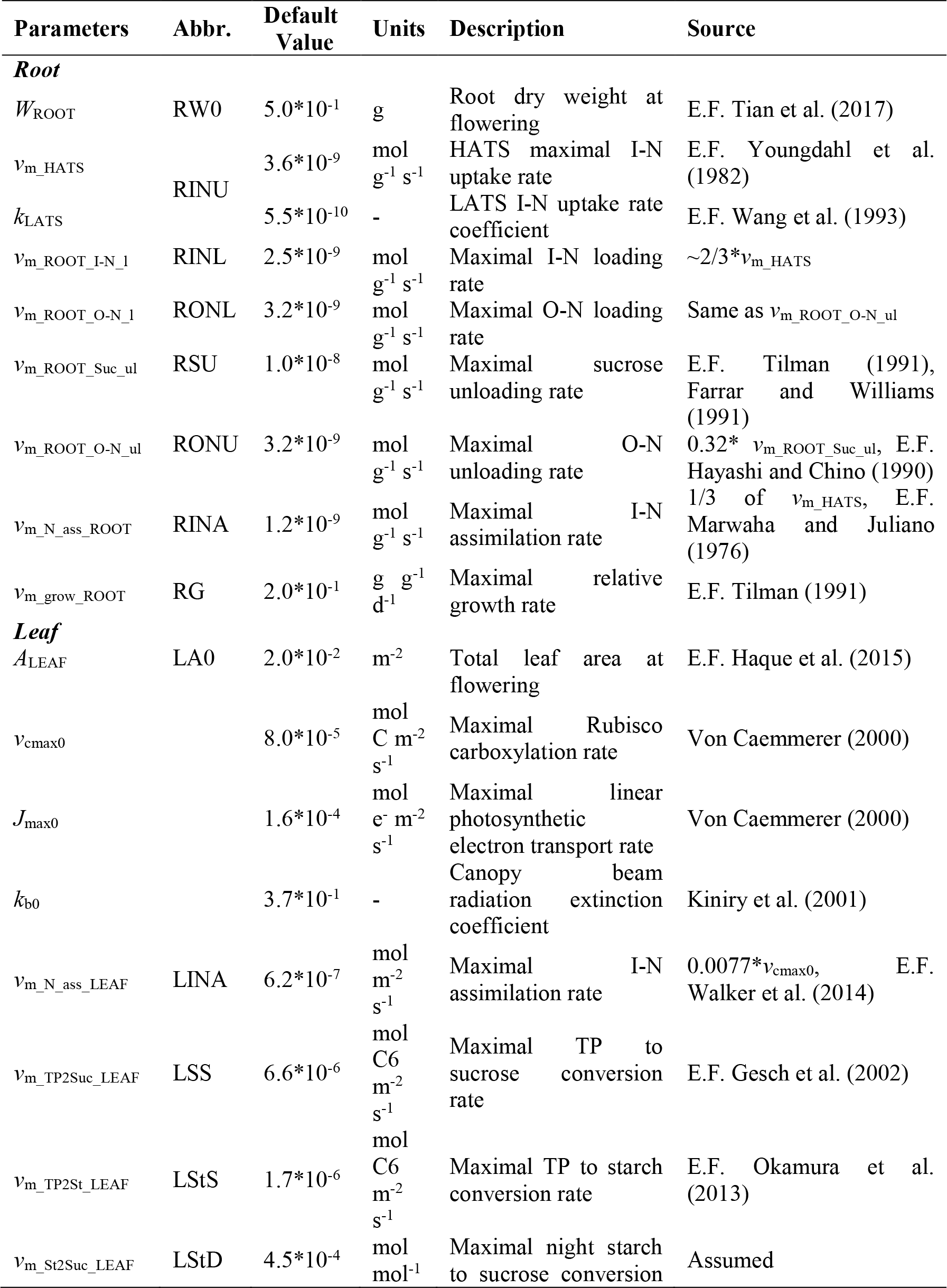
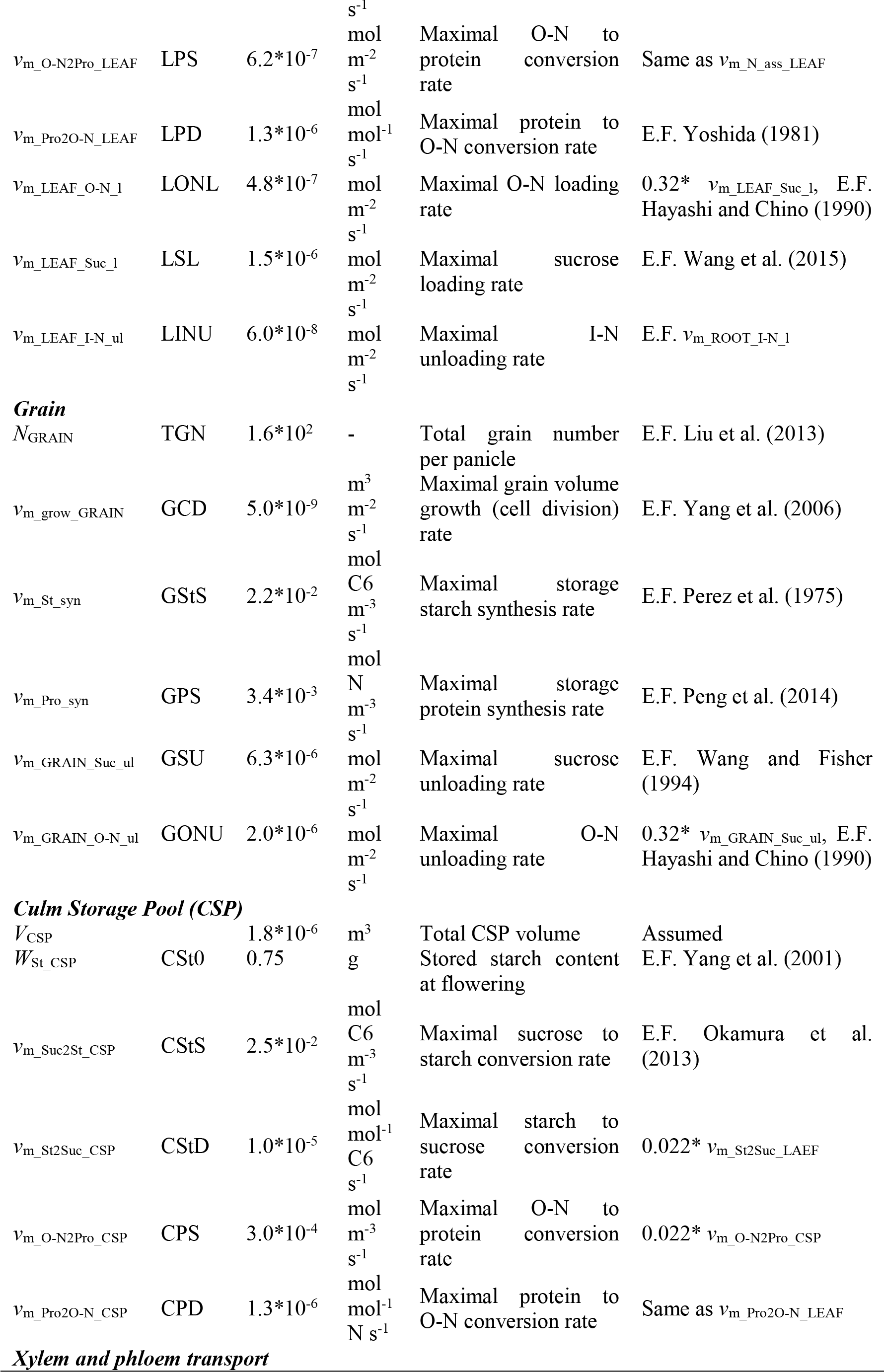
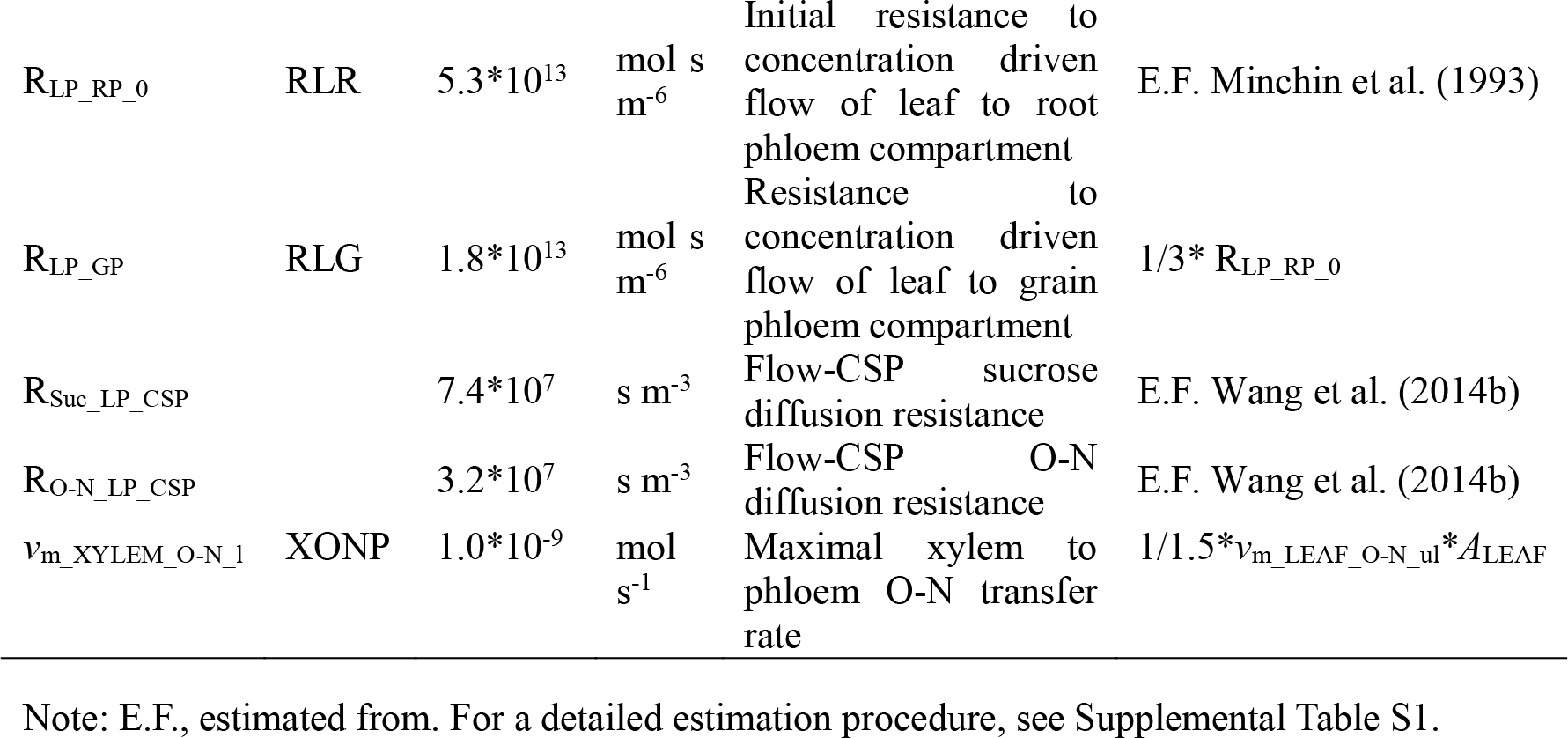
Major model parameters, their abbreviations, default values, units, descriptions and references used for parameter estimation.

To further test the performance of WACNI, we generated an *in silico* ‘natural population’ with 1000 rice individuals by randomly perturbing 32 model parameters, which comprise 28 cellular level biochemical (*v*max of enzymes and transporters) and biophysical (phloem transport resistance) parameters and 4 biomass related parameters defining plant initial state, i.e. root dry weight, total leaf area, grain number and CSP starch content at flowering. The post-flowering growth for each of these individuals was simulated. Firstly, we examined the relationship between grain yield and 4 agronomic traits which are commonly regarded being critical for grain yield. Results show a strong positive correlation between grain yield and total canopy photosynthesis for whole grain filling season (TCP, Fig. 3a) or grain filling duration from flowering to harvest (GFD, Fig. 3b), both of which have long been recognized as dominant factors controlling grain yield (Yoshida, 1981; Zhai et al., 2002; Yang et al., 2008). A decrease trend of grain yield with increase of total grain N content was predicted (GNC, Fig. 3c), which is also consistent with field observations (Triboi et al., 2006). Interestingly, although there seemed to be an increase trend for grain yield with increase in average grain filling rate, i.e. grain dry weight increase rate from flowering to harvest (GFR), a non-linear relationship was found between optimal grain yield and GFR (Fig. 3d). This explained why GFR was positively correlated with grain yield in some studies but not related to or even negatively correlated with grain yield in others (Jones et al., 1979; Yang et al., 2008).

**Figure 3.**
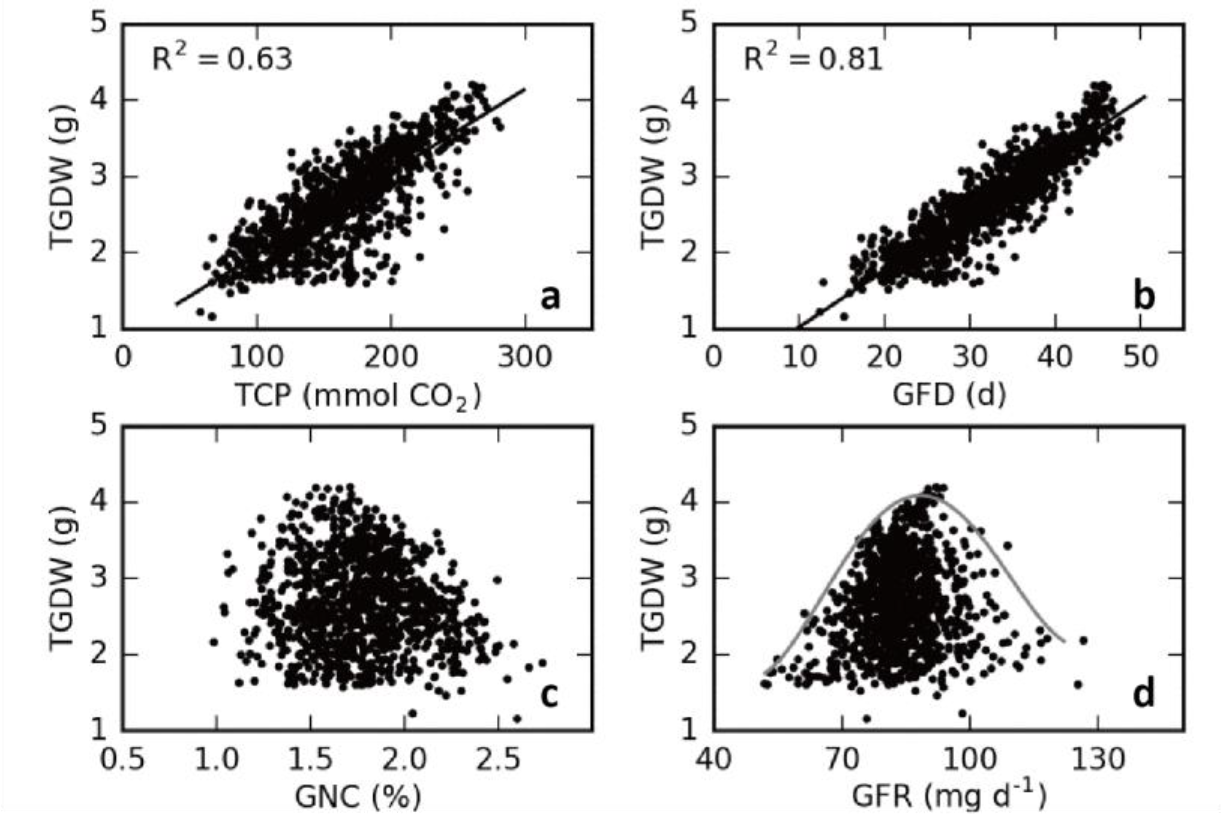
Correlation between TGDW and 4 agronomic traits within the ‘natural population’ generated by WACNI. These agronomic traits are total canopy photosynthesis for whole grain filling season (a), grain filling duration from flowering to harvest (b), final grain total nitrogen content (c) and average grain filling rate from flowering to harvest (d). The grey line in (d) illustrates the trend of optimal TGDW in the population with increasing GFR.

It’s noteworthy that predicted grain yields showed no significant correlation with any of these 32 parameters (Supplemental Fig. S1), even though each of these parameters controlls a major process involved in yield formation. This incapacity was largely due to the complex interaction patterns and covariation of large number of processes, which jointly determine crop yields. For the same reason, even if all the molecular processes were considered simultaneously, linear regression models can only maximally explained less than 50% variance of grain yields in the whole population (Supplemental Fig. S2a, b). Generally, the smaller population used to train a correlation model, the poorer predictive power (Supplemental Fig. S2b). When 100 individuals in a population were randomly selected, on a rare event, a linear model with high R^2^ might be derived (R^2^ = 0.73); however, its predictive power on the remaining 900 individuals was very poor (R^2^ = 0.09; Supplemental Fig. S3c). Overall, these results suggested a major weakness in the correlation based approach in identifying relationship between parameters and complex traits in a complex system where complex non-linear interactions between system’s components exist.

### WACNI predicted the effect of genes in mutants and/or transgenic plants

We performed an extensive literature study for genes controlling the modelled 14 biochemical and biophysical processes in rice. Ten genes were found and the impacts of genetically modifying their expressions on plant physiological traits are summarized in Table 2. Remarkably, WACNI accurately predicted most of their effects on major agronomic traits, especially on grain yields (Table 2). For example, WACNI predicted a monotonic increase of grain yield with gain-of-function of the grain starch synthesis process (‘GStS’ in Supplemental Fig. S5), which has been demonstrated in both wheat (Smidansky et al., 2002) and rice (Smidansky et al., 2003). On the contrary, WACNI predicted that when grain sucrose unloading capacity increases, the grain yield first increases then decreases (‘GSU’ in Supplemental Fig. S5). This is because when GSU is low, grain yield formation is limited by sink strength; with increase in GSU and hence increased sink capacity, plant gradually reaches its maximal grain yield as a result of more and more coordinated source and sink strengths; further increase in GSU leads to higher than optimal rate of C and N utilization in the grain sink, which leads to decreased accumulated canopy photosynthesis and hence grain yield (‘GSU’ in Supplemental Fig. S5). These predictions are consistent with results of the transgenic study on the effect of changing the strength of a grain sucrose transporter *GIF1* (Wang et al., 2015). In addition, WACNI predicted a strong negative impact of impaired function of leaf sucrose loading (LSL) on grain filling rate, total canopy photosynthesis, harvest index and final grain yield. Although grain yield does not decrease with increasing LSL capacity, WACNI predicted a leaf sucrose loading capacity (LSL) higher than 0.6-fold of the default value can no longer increase grain yield in CK plant (‘LSL’ in Supplemental Fig. S3). Consistently, Eom et al. (2011) found that impaired function of *OsSUT2*, which exports sugar from leaf, led to severe growth retardation and yield decrease, whereas overexpression of *OsSUT2* in the mutant plants exhibited no more gain of yield compared to CK plants. WACNI also predicted a minor impacts of altering leaf starch synthesis (LStS), leaf sucrose synthesis (LSS) and culm starch synthesis capacity (CStS) on grain yield, which again is consistent with earlier experiments (Table 2).

**Table 2.**
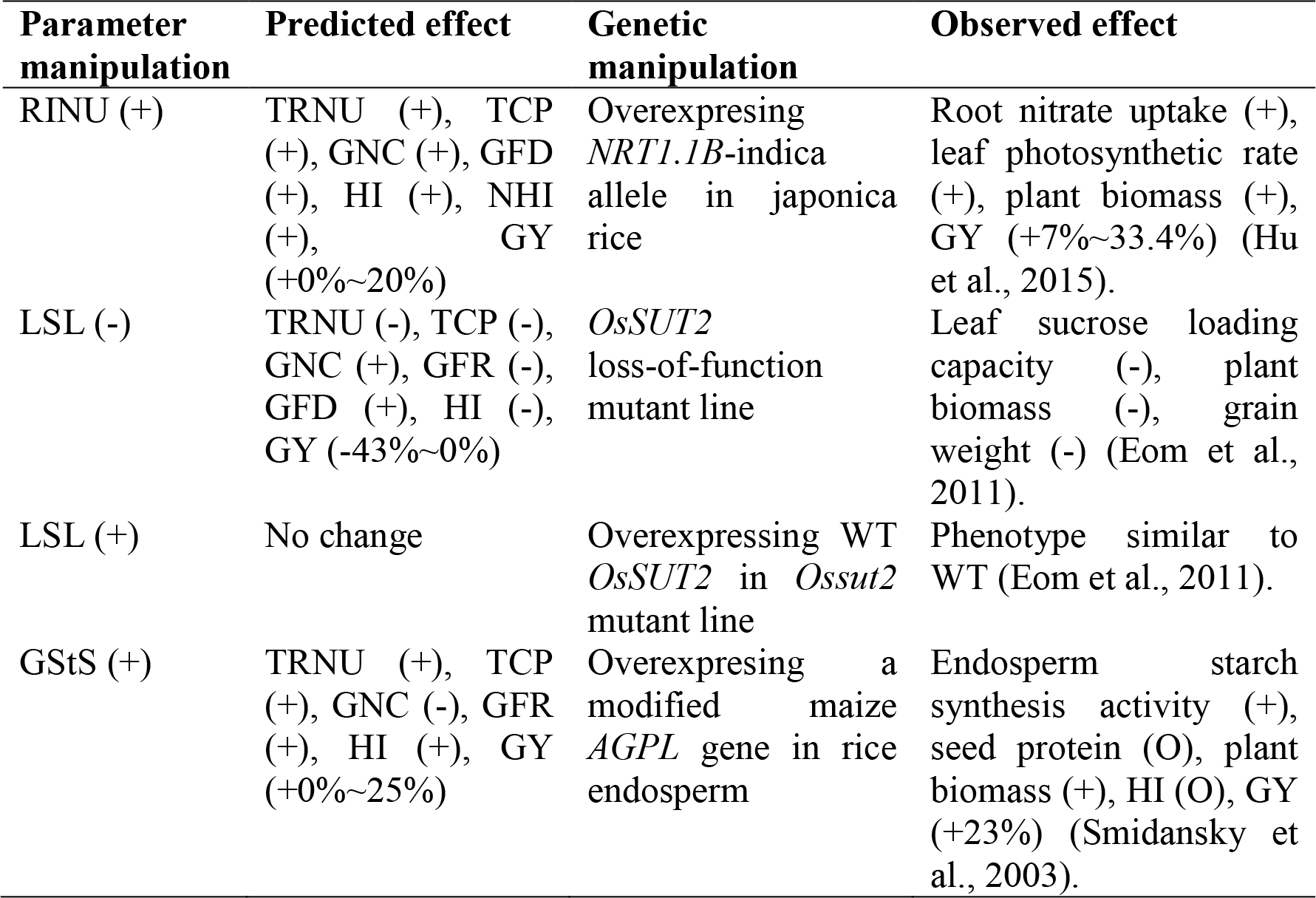
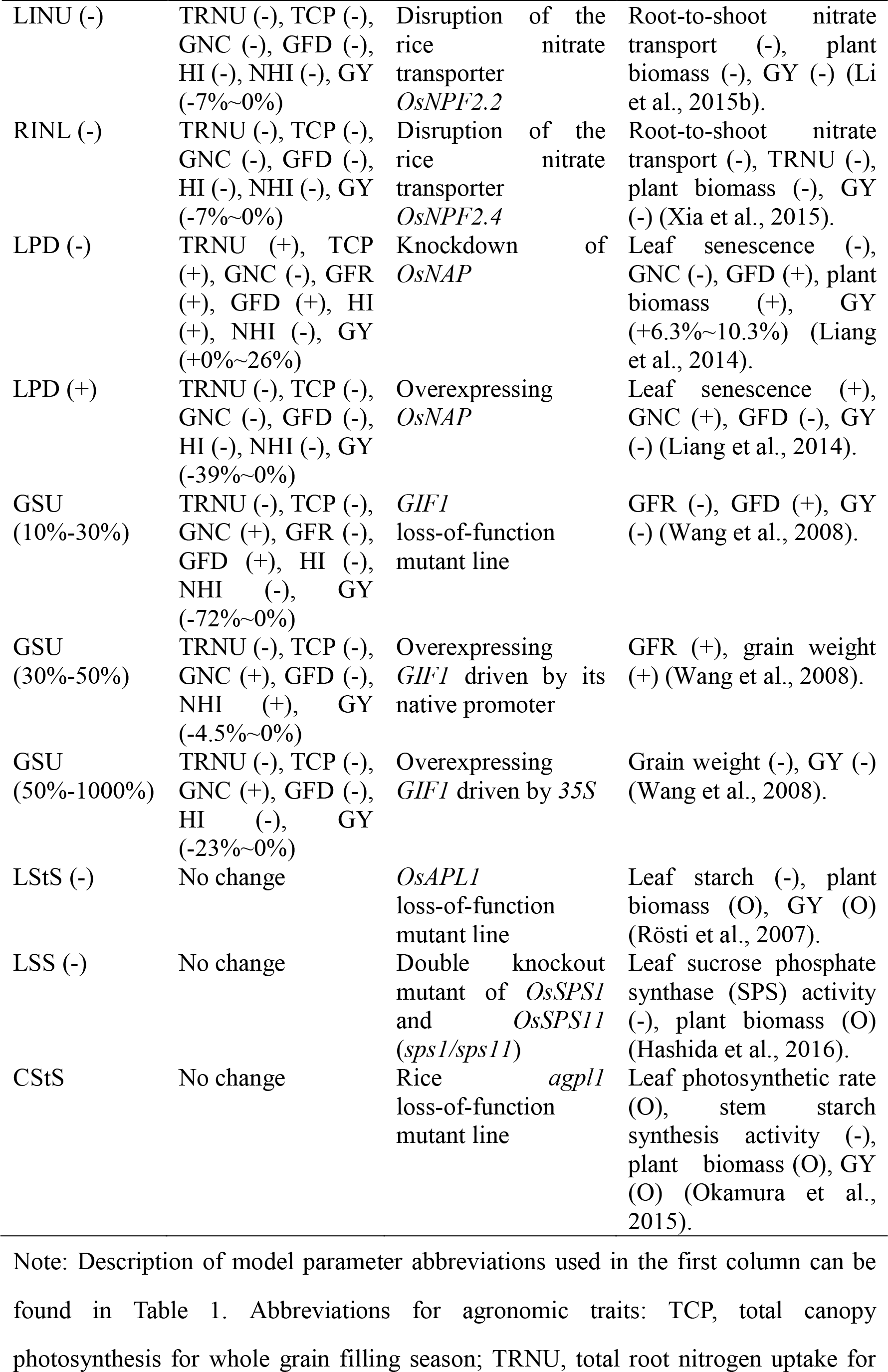
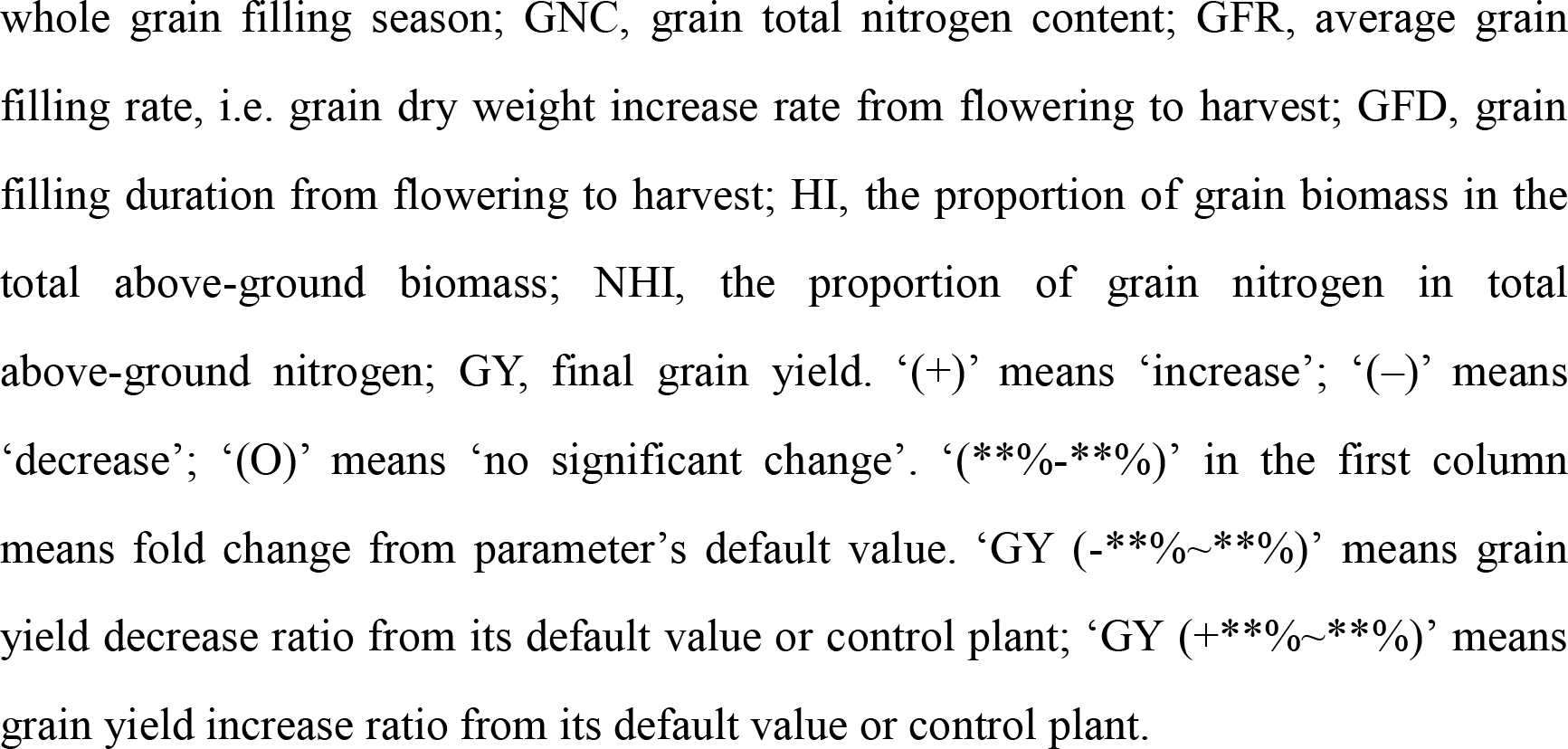
Predicted and observed response patterns of grain yield and other agronomic traits to genetic manipulations of genes controlling one of the modelled 14 molecular processes. The predicted effects of these parameters on all 8 agronomic traits were presented in Supplemental Fig. S3-9.

### Classification of molecular modules influencing grain yields

With WACNI, we first systematically explored the influence of all the above mentioned 28 cellular parameters on grain yield and other agronomic traits (Supplemental Fig. S3-9). These agronomic traits include the above mentioned TCP, GFD, GNC, GFR, and total root nitrogen uptake for whole grain filling season (TRNU), above-ground dry weight harvest index (HI) and above-ground nitrogen harvest index (NHI). Each of the 28 parameters, which is involved in one of C and N metabolism and transport processes, reflects the function of either one or a set of genes forming a molecular module. Based on the predicted patterns of modifying these parameters on grain yields, we classified these parameters into four types (Table 3). Specifically, root N uptake capacity exemplifies a universal yield enhancer (UYE), which monotonically enhances grain yield by increasing TCP and HI (Fig. 4a-d). In contrast, universal yield inhibitors (UYIs) can monotonically inhibit grain yield, which is exemplified by leaf protein degradation capacity (Fig. 4e-h). Conditional yield enhancers (CYEs) can non-linearly influence grain yield. For example, increasing grain cell division capacity can first increase then decrease grain yield as a consequence of its negative effect on TCP, its positive effect on GFR and its curvilinear effect on HI (Fig. 4i-l). Finally, some molecular modules show little or no influence on grain yield and hence termed as weak yield regulator (WYR, Fig. 4m-p). In spite of 10 parameters for which the impacts of their genetic manipulations had been studied in rice as mentioned above, WACNI predicted several new targets which may dramatically influence grain yield. For example, increasing culm stored protein degradation capacity (CPD) can benefit grain yield, whereas increasing leaf amino acids export capacity (LONL) after flowering may reduce grain yield (Table 3). These become natural targets for future experimental testing.

**Figure 4.**
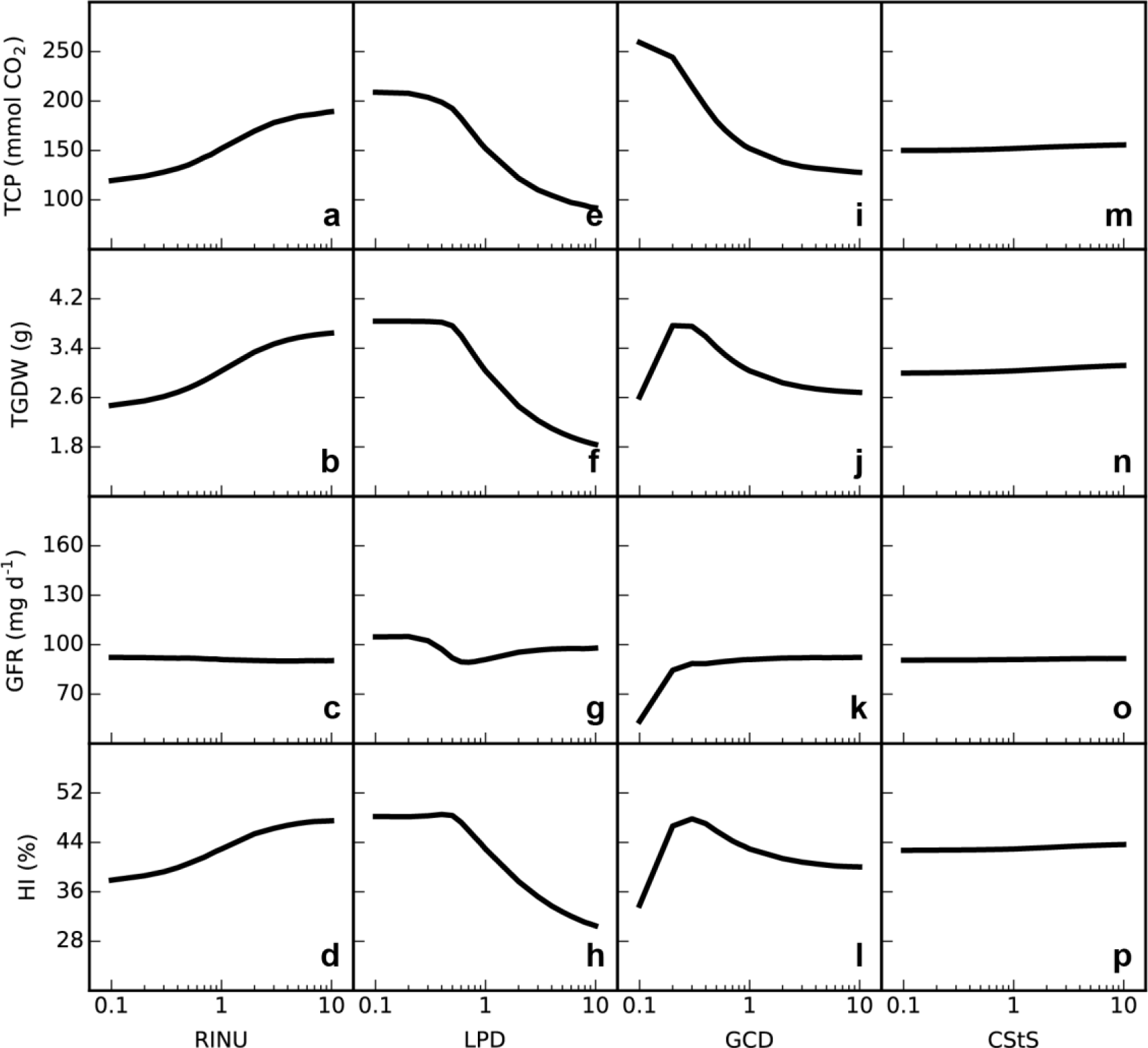
The simulated impacts of changing single model parameter (fold change from default value) on major agronomic traits. The parameters are root I-N uptake capacity (RINU), leaf protein degradation capacity (LPD), grain cell division capacity (GCD) and culm starch synthesis capacity (CStS). The agronomic traits are total canopy photosynthesis for whole grain filling season (TCP), total dry weight of all grains in the panicle (TGDW), average grain filling rate, i.e. grain dry weight increase rate, from flowering to harvest (GFR) and above-ground dry weight harvest index (HI). Note: the x-axes are log-scaled.

**Table 3.**
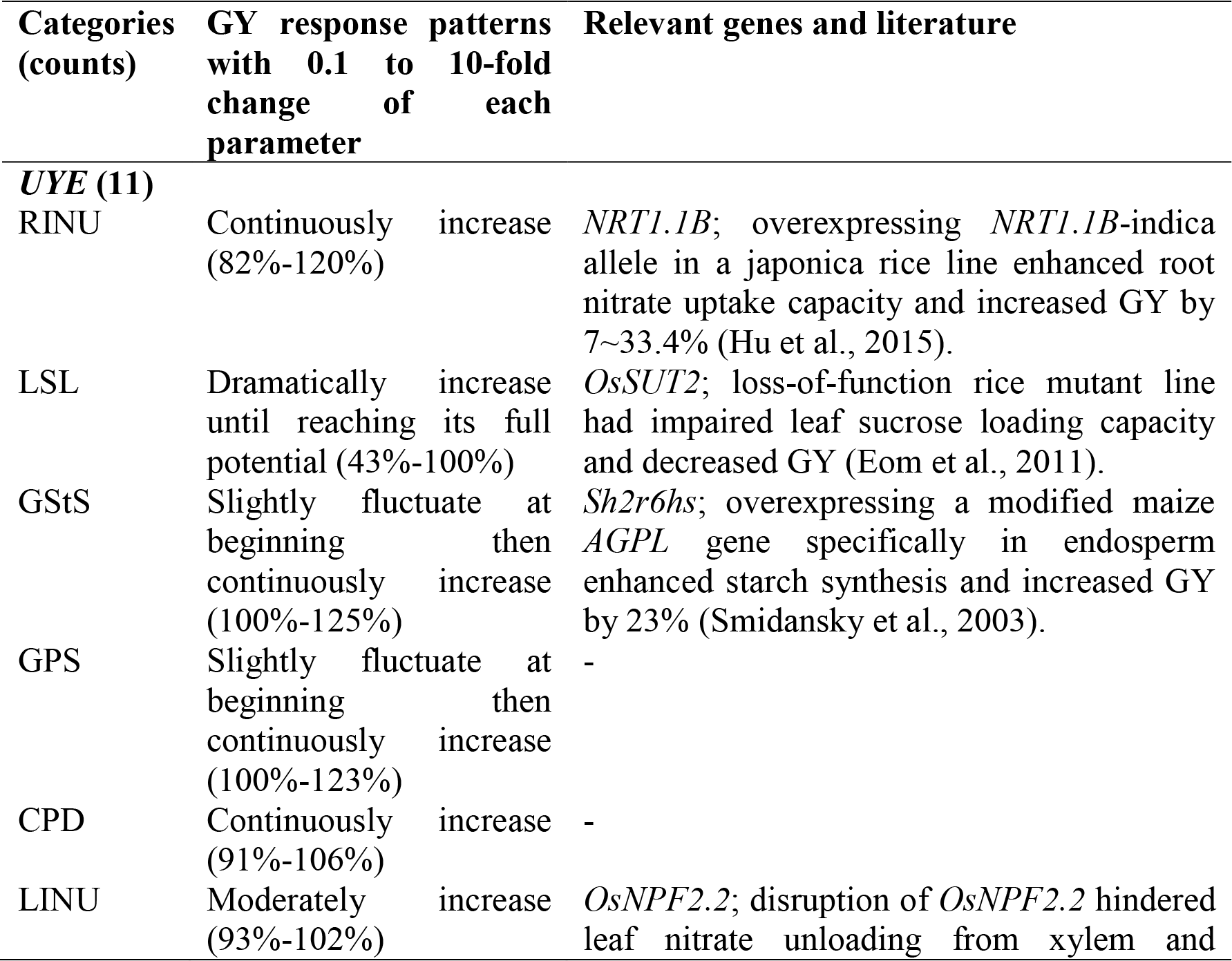
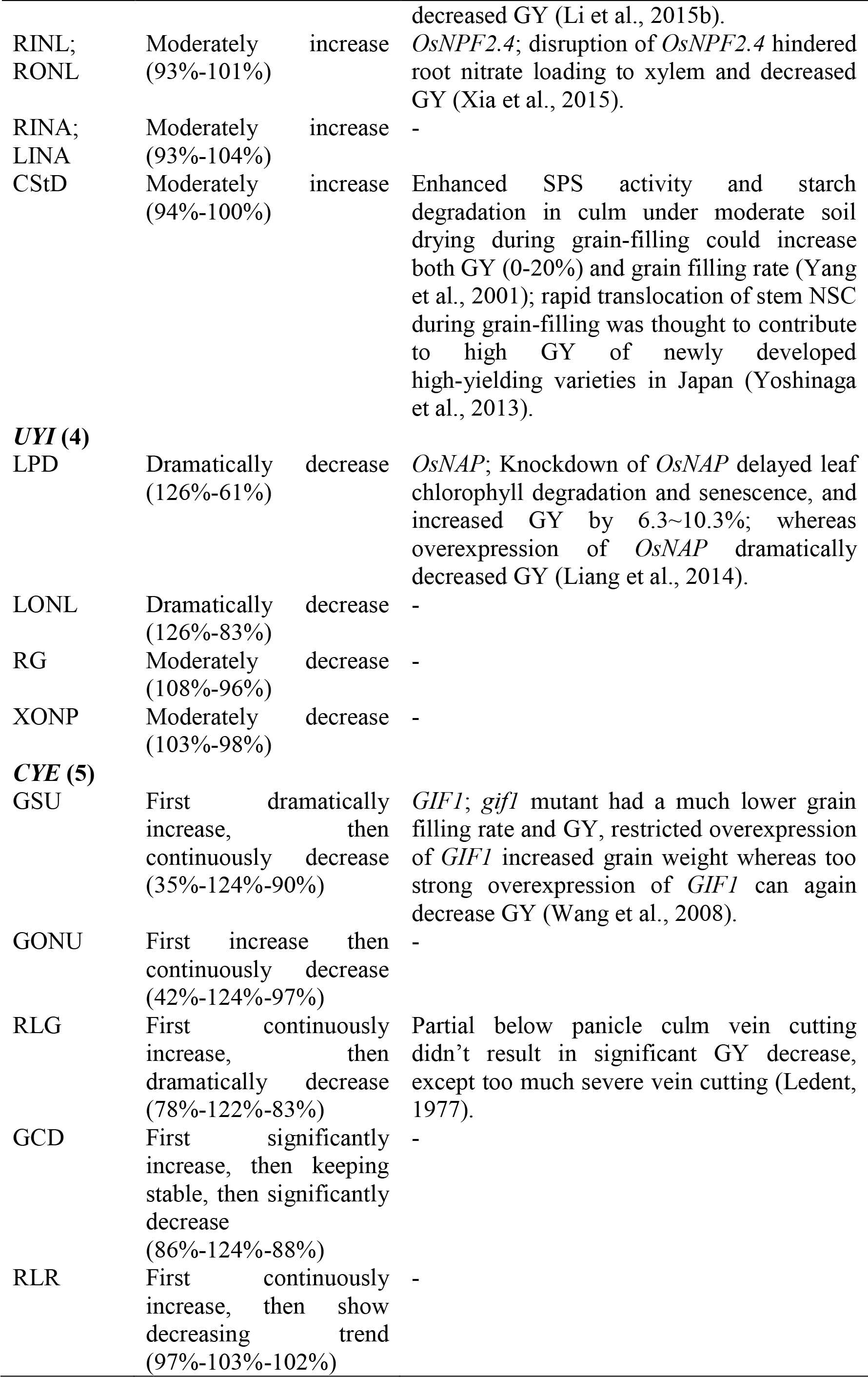
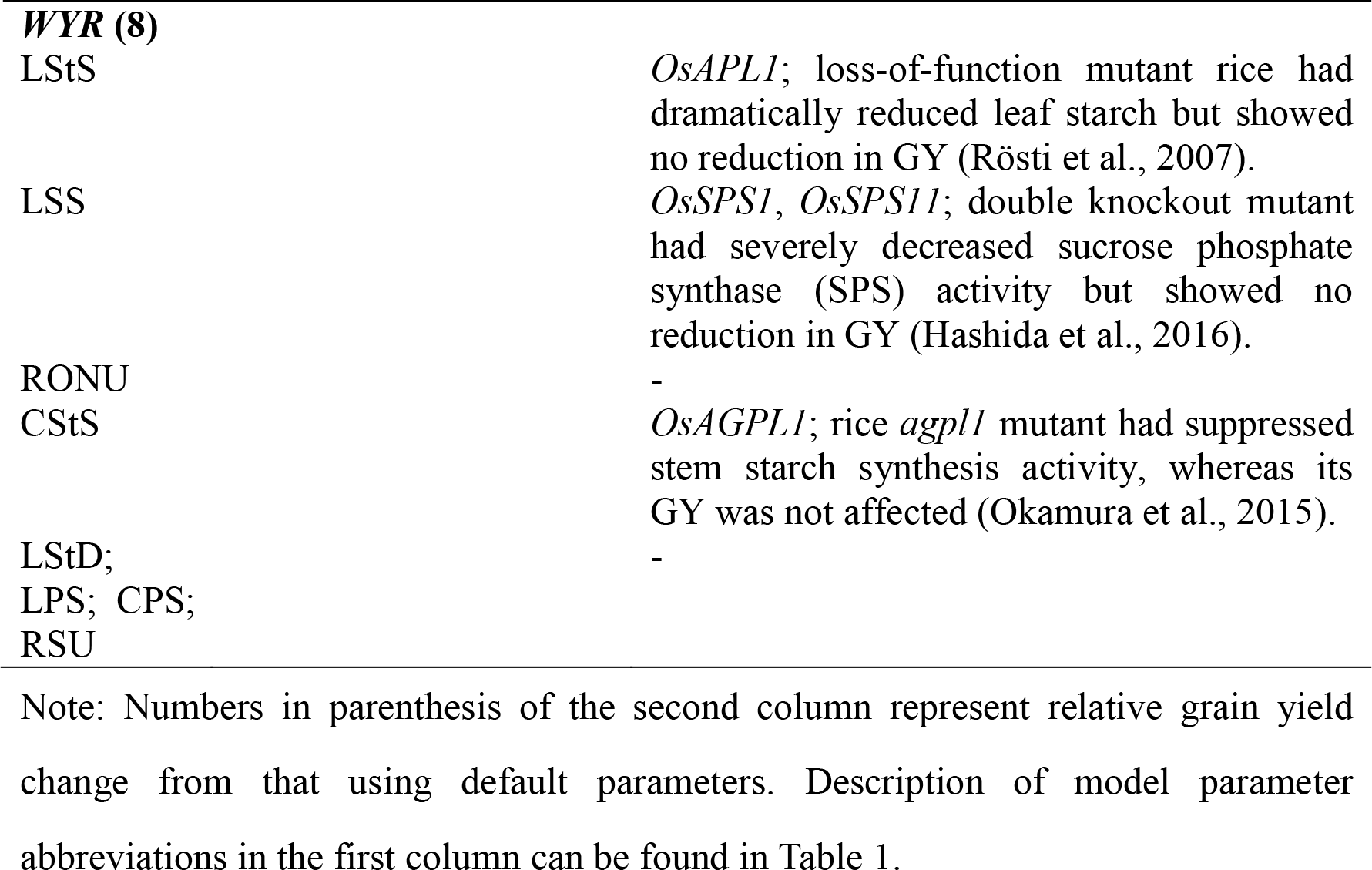
Classification of the biochemical/biophysical parameters based on their impacts on grain yield (GY). See more detailed response patterns of yield and other agronomic traits to change of each parameter in Fig. S3-9.

### Molecular and macroscopic physiological features of high-yield rice under the current condition

We further used WACNI to identify molecular and macroscopic physiological features associated with high-yield rice lines. To do this, we used WACNI to evolve *in silico* five independent populations by varying 32 parameters mentioned above with high grain yield as the selection target. At the end of the *in silico* evolution, we found that the highest grain yields in these five populations were similar (Supplemental Fig. S10), suggesting that the model reaches a global optimal solution. The top 1% individuals which had the highest final grain yields in each population were then merged to form an elite group. Result showed that values of some parameters, e.g., LPD, LONL, GStS, RINU and RG, were relatively conserved in the elite group, i.e. they showed small variations (Fig. 5a). On the contrary, though individuals in the elite group all have similar high grain yields (‘Elite’ in Fig. 5b), up to 4-fold differences in the parameter values exist for a number of molecular level parameters, e.g., LINA, LSS, CStS, and RONL (Fig. 5a). When the maximal and minimal values of each parameter in the elite group were used to generate two new individuals, named as ‘Max’ and ‘Min’ (Fig. 5b), respectively, we found that both of them have much lower grain yields than the average grain yield in the elite group (‘Elite’ in Fig. 5b), suggesting the necessity of the systems design to identify optimal molecular-module combinations for high yield.

**Figure 5.**
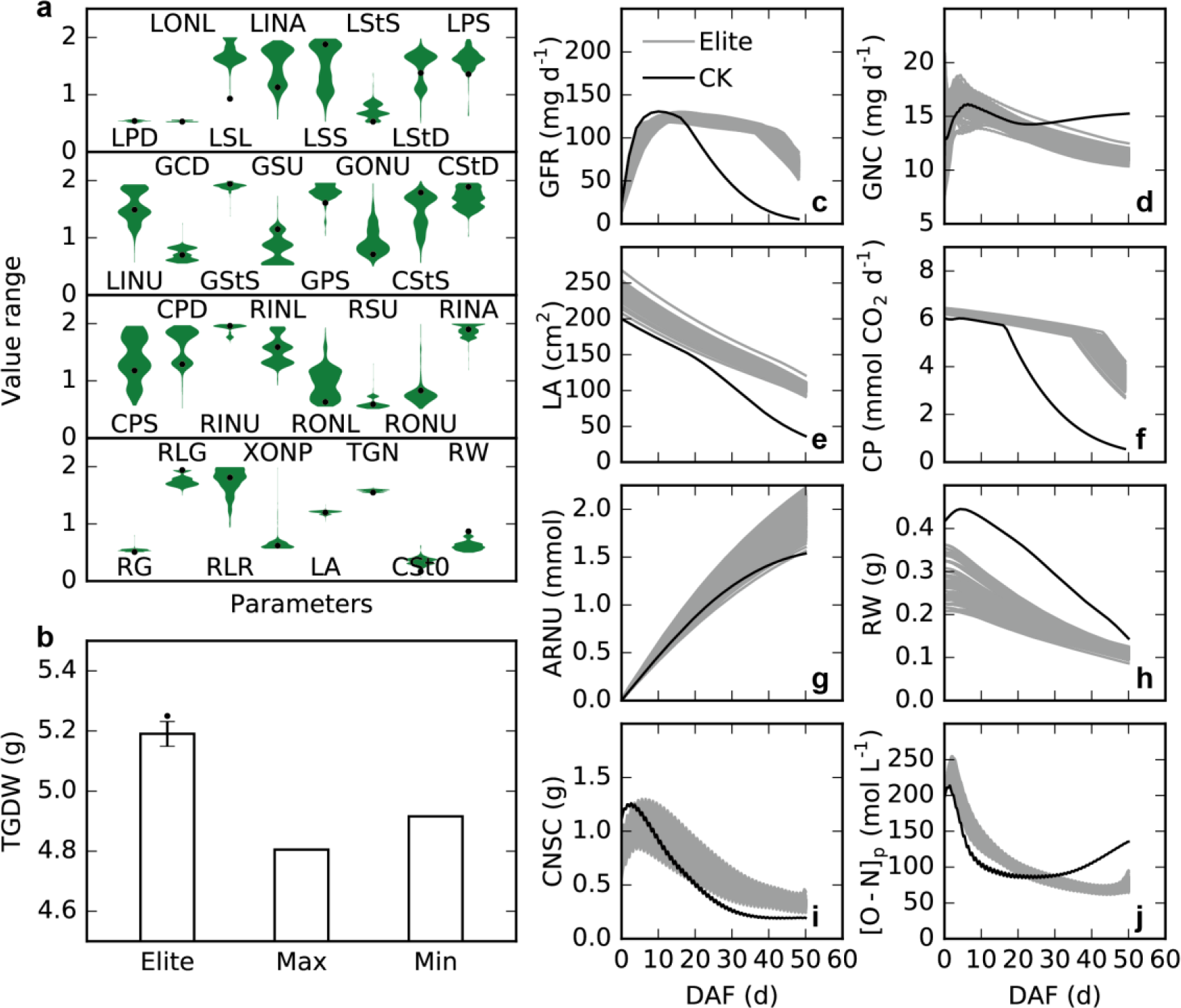
Molecular and macroscopic physiological features of high-yield rice under normal condition. **a**, distribution of relative value of each model parameter (fold change from default value) for *in silico* high-yield rice lines in high-yield elite group. **b**, comparison of average grain yield of elite group (n=500, ‘Elite’) and grain yield of individuals with parameters assigning to maximal (‘Max’) or minimal (‘Min’) values in elite group, respectively. The black dots in **a** and **b** show the parameter values and grain yield of the highest yielding rice in the group, respectively. **c-j**, comparison of macroscopic physiological traits of high-yield rice lines in elite group and CK rice line. Abbreviations: TGDW, total grain dry weight; GNC, grain total N concentration; LA, total leaf area; CP, canopy photosynthesis; TRNU, total root I-N uptake; RW, root dry weight; CNSC, culm non-structural carbohydrates; [O-N]_p_, phloem O-N concentration.

In contrast to large variations among molecular level parameters in the elite group, macroscopic physiological features during grain filling for the elite group were more conserved (Fig. 5c-j). Furthermore, the elite group, as compared to the CK individual which uses default parameter values, shows distinct dynamic pattern of macroscopic physiological features along grain filling. Firstly, in the elite group, the grain filling rate increased slower at the early grain filling period but kept a high level for much longer duration (Fig. 5c); the final grain N concentration was lower (Fig. 5d). Secondly, the elite group showed slower rate of decrease in both leaf area and canopy photosynthesis rate at the late grain filling period (Fig. 5e-f), a feature commonly termed as ‘stay green’ (Jagadish et al., 2015). Surprisingly, the elite group had a smaller root mass at flowering (Fig. 5h) but higher accumulated N uptake (Fig. 5g) as a result of its more efficient N uptake capacity (RINU, Fig. 5a). Finally, the elite group had less non-structural carbohydrates storage (CNSC) at flowering but higher CNSC during the late grain filling period (Fig. 5i); the O-N in phloem ([O-N]_p_) showed a steady decline in the elite group, in contrast to the CK where [O-N]_p_ gradually increased at the late grain filling period (Fig. 5j).

### Optimal C-N interaction and underlying molecular features for high yield under changed environmental conditions

The same evolutionary strategy was used to identify molecular and macroscopic features associated with high grain yield under different environments, i.e. high ambient CO_2_ (HC), low light (LL) and low soil N (LN), respectively. We examined values of the twelve most conserved parameters identified under normal condition (NC; Fig. 5a) for the top 1% high-yield individuals in each condition. Again, on one hand, several parameters showed small variations, i.e., LPD, LONL, GStS and RINU (Fig. 6a), indicating their essential and conserved roles in regulating grain yield under a number of conditions. On the other hand, some parameters showed high levels of environmental dependency. For example, for the purpose of achieving high grain yield, a substantial increase in grain cell division rate (GCD) under HC, but the opposite under LL and LN compared to that in NC, was predicted to be critical (Fig. 6a).

**Figure 6.**
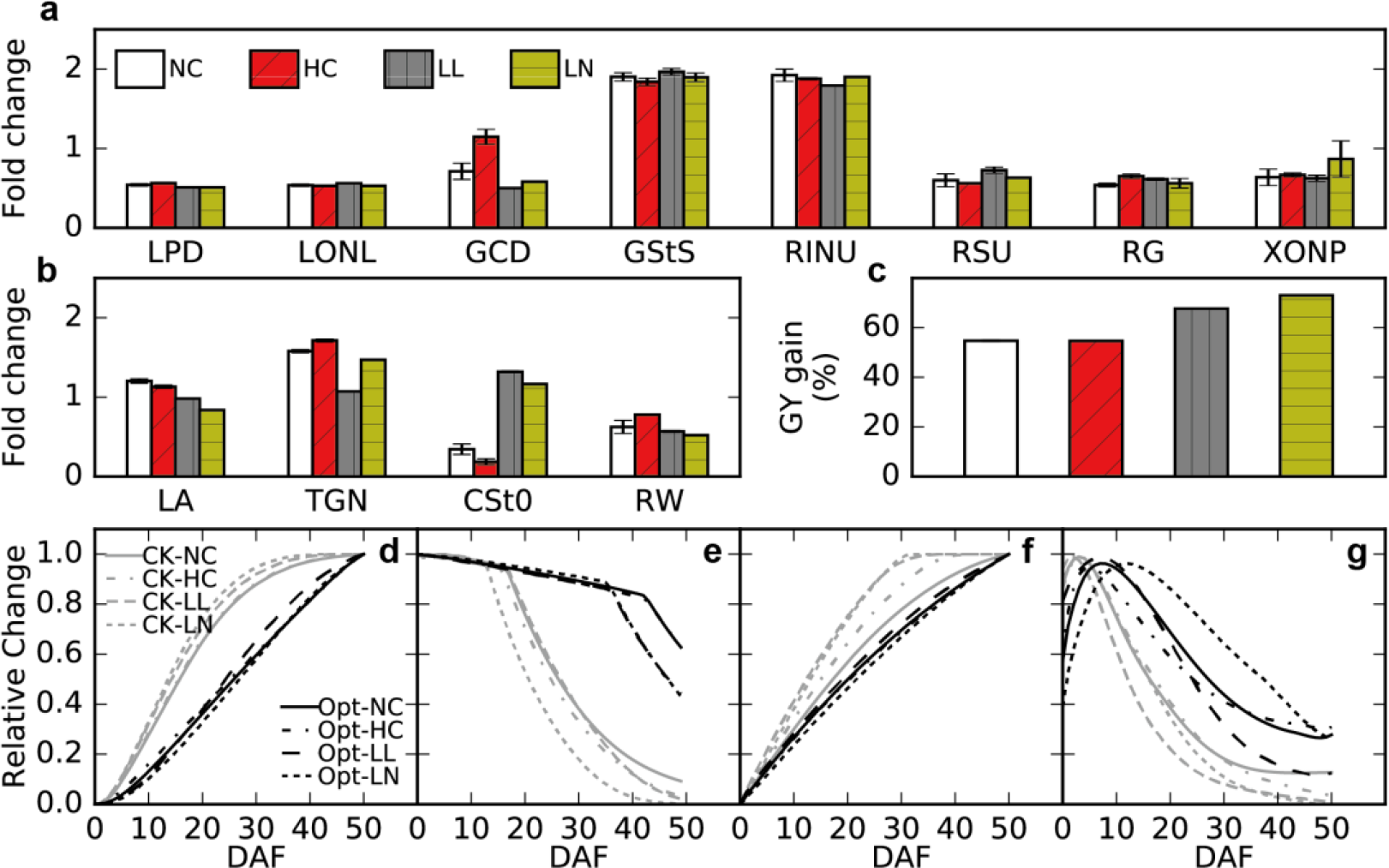
Molecular and macroscopic physiological features of high-yield rice under different environments. Optimal combinations of 8 biochemical parameters (**a**) and 4 biomass parameters (**b**) for high yield under various environmental conditions. The y-axis shows fold change of each parameter from its default value. **c**, grain yield gain of highest yielding lines compared to CK plants under different environmental conditions. **d-g**, comparison of normalized plant physiological characteristics for highest yielding and CK rice plants under different environmental conditions. **d**, total grain dry weight; **e**, canopy photosynthesis rate; **f**, total root I-N uptake; **g**, culm non-structural carbohydrates storage. The four environmental conditions are normal (NC), high ambient CO_2_ (HC), low light (LL) and low soil N (LN).

Several key physiological features common to all high-yield rice under different environments were also identified. These features included a sustained and steady growth of grain sink (Fig. 6d), a prolonged leaf photosynthetically active period (Fig. 6e), an efficient rather than large root system and a sustained root N uptake (Supplemental Fig. S11g; Fig. 6f), and more CSP carbohydrates available for late phase grain filling (Fig. 6g). Although the values of these features differed (Supplemental Fig. S11b, e, f, h), their relative changes during the grain filling period largely remained the same throughout all different conditions (Fig. 6d-g), suggesting that these patterns or growth strategies are necessary macroscopic features and can be used as selection targets in the high-yield rice breeding programs. In particular, WACNI predicted a 54.6% to 73% grain yield enhancement compared to CK plants under different scenarios when pyramiding all these superior features properly with a systematic rational design (Fig. 6c).

## DISCUSSION

### WACNI: a living module mechanistically describing plant growth based on molecular processes

Develop a complete *in silico* representation of plant growth and developmental process is a major challenge facing plant biologists (Zhu et al., 2015; Marshall-Colon et al., 2016; Xiao et al., 2017). Such a model needs to include not only the basic metabolic processes in different organs of a plant, source sink interactions, and also the plant environment interactions (Zhu et al., 2015; Xiao et al., 2017). On this aspect, many component modules for different plant processes, such as leaf photosynthesis, stomata conductance, flowering date determination, root morphogenesis and nutrient update, above ground morphogeneis, canopy photosynthesis, canopy atmosphere interaction, etc., which are all needed to develop such a complete model, have been developed, see detailed reviews by Zhu et al., (2015). What is largely missing now is a highly mechanistic model of plant growth and development capable of predicting the non-linear interactions between processes in different plant tissues.. Here we report the development of the Whole plant Carbon Nitrogen Interaction model (WACNI) to fill in this gap. By faithfully describing the detailed biochemical and biophysical processes in different organs, WACNI accurately predicts the dynamic changes in different biochemical and physiological processes during the grain filling stage (Fig. 2) and successfully predicted the currently reported impacts of genetic manipulations on grain yields (Fig. 4; Supplemental Fig. S3-9; Table 2; Table 3). As its initial application, WACNI was used to identify the major biochemical and macroscopic features associated with high yield (Fig. 5-6). It is worth emphasizing here again that completely different from all previous models of crop source sink interaction which are mainly based on either preset rules on the relationship between source and sink tissues, or empirical relationship derived from measured data, WACNI predicted source sink interaction as an emergent property from molecular mechanisms in different tissues. This feature enables the model to directly link macroscopic or physiological features to molecular processes, hence provides a direct bridge between crop models and current crop genetics research.

To ease its easy reuse or extension, WACNI follows a principle of modular design. In WACNI, the metabolic processes involved in leaf, grain, root, and CSP were described separately as individual modules, which hence can be replaced by other modules or extended by other researchers easily. As a result, WACNI can be used as a ‘living’ system, i.e. it will be continuously updated or improved by incorporating more and more detailed molecular mechanisms with time. In addition, most of processes described in WACNI are rather generic among crops or plants; so WACNI can be easily adapted and parameterized for other crops or plants, in particular for other cereal crops. WACNI hence can be used as a new basis or chasis for different crop growth and development models.

### WACNI: a theoretical framework to study whole plant carbon nitrogen interaction and crop yield formation

With detailed consideration of C and N metabolic reactions and the major feedback regulatory mechanisms in different tissues, WACNI provides a tool to study the metabolic interactions between source and sink organs and how these interactions influence crop growth and yield. Currently, the regulatory mechanisms incorporated in WACNI include stimulation of N on leaf photosynthesis and leaf longevity, inhibition of carbohydrate on leaf photosynthesis, stimulation of carbohydrate on root growth and I-N uptake, and inhibition of N on root I-N uptake (see Supplemental Data File S1). As a result of these interactions, WACNI predicted that when a particular gene is altered, even only in one tissue and at one particular growth stage, a multitude of changes could occur in different processes in different organs

(Supplemental Fig. S3-9), which is a phenomenon that had been observed for long (Wilson, 1988). WACNI can simulate these different processes simultaneously and therefore can help study the sequence of actions or effects after a particular molecular process is altered by systematically comparing the predicted and observed changes at molecular, physiological and yield levels.

Similarly, WACNI can be used to study the non-linear response of yield to manipulation of a particular gene. This is important since most genes influencing source, sink and transport processes identified so far (Takeda et al., 2001; Xing and Zhang, 2010; Long et al., 2015) show pleiotropic impacts on yields for different cultivars and/or under different growth conditions (Chang et al., 2017). Why is this so? First, grain yield, as a complex trait, is controlled by many factors simultaneously; the status of these different factors jointly determines the potential impact of manipulating a particular factor on the crop yield. Thus, the variations of protein abundance or enzyme activities in different cultivars will inevitably lead to different responses in yield when a particular process is manipulated. For example, WACNI predicted a leaf sucrose loading capacity (LSL) higher than 0.6-fold of the default value can no longer increase grain yield in CK plant (LSL in Supplemental Fig. S3), whereas a typically 1.5-fold LSL was observed for high-yield lines (LSL in Supplemental Fig. S11a). Secondly, grain yield might have a highly non-linear and non-intuitive responses to modification of a gene. For example, WACNI predicted that when grain sucrose unloading capacity (GSU) was increased, the grain yield first increased then decreased (Supplemental Fig. S5; Wang et al., 2015). Simulations using WACNI suggest that besides these 4 biomass related parameters (LA, TGN, CSt0, RW; Fig. 6b), grain organic N unloading capacity and cell division rate (GONU and GCD), and biophysical resistance of long distance phloem transport (RLG and RLR) (Table3; Supplemental Fig. S5, 6, 9) all influence grain yield in a non-linear manner, suggesting that all these need to be fine-tuned for higher grain yield.

### WACNI: a guide for the future synthetic biology effort of engineering crops for greater yield

The ability of WACNI to faithfully predict the non-linear interaction between source and sink tissues promises its application in guiding current efforts of engineering or breeding crops for desired traits. In most contemporary crop breeding programs, features used in crop breeding are based on either breeders’ experience or results from correlation studies using high-yield lines. Although being highly successful historically, these approaches become less effective recently as indicated by the stagnancy in the enhancement of yield on a land area basis in the last decade (Ray et al., 2012). This stagnancy has been commonly attributed to a number of factors, including early senescence of root and leaf (Liang et al., 2014), inefficient remobilization of stem and sheath storage material (Yang et al., 2002), and the dramatically weakened grain-filling of the inferior grains in the late grain filling period (Yang and Zhang, 2010). Though there are efforts in manipulating these individual features, analysis based on WACNI suggests that these can all be regarded as syndromes of a suboptimal whole-plant level C-N interaction. In fact, WACNI suggests that the degree of sustained and steady growth of grain sink can be used as ‘physiological indicator’ to evaluate degree of optimality of the whole-plant C-N interaction (Fig. 6d). Based on this, WACNI was used to design a number of molecular engineering and pyramiding schemes to gain the promised high grain yield (Fig. 5a; Fig. 6a, b; Supplemental Fig. S11a). Out of the identified engineering options, the genes that confer monotonic impacts on crop yield, i.e. the UYEs and UYIs, can be used as the first set of genes to test, followed by those genes that can conditionally influencing crop yield (CYEs), where precise control of gene expression levels (Rodriguez-Leal et al., 2017) is needed to gain the desired engineering output.

## CONCLUSION

This article presents a whole-plant level mechanistic model of carbon-nitrogen interaction, WACNI, which bridges complex molecular processes with macroscopic physiological properties and crop yield. Considering that systems model guided rational engineering of photosynthesis has recently gained huge success (Kromdijk et al., 2016), the WACNI model presented here effectively can support systems model guided rational engineering of crops for greater yields. We envisage that WACNI, together with the rapid advances in genome editing technologies and gene cloning, powered by the state of art genomics and phenomics capacities, lays the foundation for an era of model guided crop molecular breeding by design.

## MATERIALS AND METHODS

### Model construction and parameterization

Fourteen different types of primary biochemical/biophysical processes were incorporated in WACNI (Fig. 1). The rate equations are described in four different subsections according to the organs they present in, i.e. root, leaf, grains and culm in supplemental materials (Eqn 1.1-14.1 in Supplemental Data File S1). Respiration for each organ is described in the subsequent *Respiration* subsection (Eqn 15.1-15.4 in Supplemental Data File S1). Supplemental Table S1 provided description of parameters used in equations. The major hypotheses used during model construction are presented in Supplemental Data File S3.

The kinetic parameters for processes modeled in WACNI are derived from a vast amount of genetic, molecular biochemical and plant physiological research in rice or other cereal crops. Michaelis-constant (*K*_m_) and substrates feedback regulatory factor (*K*_e_) for transport processes are estimated based on organ metabolites concentrations. A detailed description of parameters, their values, and the methods used to derive or estimate their values are given in Supplemental Table S1.

### Model validation

WACNI was validated by comparing the predicted and measured (from literature) physiological dynamics in different organs during the grain filling period, including organ growth and senescence, gas exchange and metabolites concentrations (Fig. 2). The accuracy of WACNI was further evaluated by reproducing statistical observations in natural rice population and predict single parameter perturbations

To conduct parameter perturbations, the *v*max in each process and two phloem resistance parameters (in total 28 parameters) were perturbed within [0.1, 10] folds to their default values individually in the model to calculate grain yield and value of other agronomic traits. These agronomic traits include total canopy photosynthesis (TCP) and total root nitrogen uptake (TRNU) for whole grain-filling season, grain total nitrogen content (GNC), grain filling duration (GFD) and average grain filling rate (GFR) from flowering to harvest, dry matter harvest index (HI) and nitrogen harvest index (NHI). The GFD was determined as the time interval between flowering day and the day when total grain dry weight (TGDW1) reaches 95% of its maximal value at the end of the simulation, and GFR was calculated as:

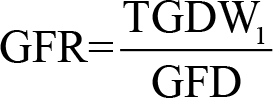

To the reproduction of statistical observations in natural rice population, the above mentioned 28 molecular biochemical/biophysical parameters and an additional four organ biomass related parameters, i.e. root dry weight, total leaf area, grain number and culm starch content at flowering, were perturbed independently for 1000 times by randomly multiplying a scaling coefficient ranging from 0.5 to 2 to generate an in silico ‘natural population’ with 1000 virtual rice lines. For the sake of comparison between lines, we set the total energy stored in biomass at flowering to be the same, i.e. when more root/leaves/grains were grown, less culm starch would be stored. The energy content of root, leaf, stem and grain are RW0/*η*_grow_, LA0*SLW/*η*_grow_, CSt0, TGN*HW/*η*_grow_, respectively, in which *η*_grow_ is growth efficiency, SLW is specific leaf weight, HW is husk weight per spikelet.

A step-wise linear regression (Draper and Smith, 2014) was used to derive relationship between grain yield versus all 32 parameters, where parameters were added into the linear model one by one based on their significance *p*-values for correlation with grain yield.

### Identification of optimal parameter combinations to maximize grain yield under different environmental conditions

A genetic algorithm was used to identify the optimal parameter combinations to maximize grain yield. In brief, this algorithm mimics the process of natural biological evolution. Here grain yield is used as the selection pressure in the algorithm.

To avoid a local optimum at the end of the evolution, five independent evolutionary populations under normal condition (ambient CO_2_ concentration is 400 µbar; soil I-N concentration is 0.14 mol m^-3^; grain-filling season is 50 days; see light intensity, air temperature and humidity information in Supplemental Fig. S12) were constructed. One evolutionary population was constructed for each of the four different conditions, i.e. high ambient CO_2_ (700 μbar), low light (half of that in normal condition) and low soil N (one tenth of that in normal condition). Each of the above 8 evolutionary populations has 100*100 individuals, i.e. 100 generations with 100 individuals in each generation. Each parameter was restricted to vary between 0.5 to 2-fold of its default value and the total available energy at flowering was set to be the same for all individuals as what had been described in the construction of ‘natural population’.

## Author contributions

T.C. performed the analysis; X.Z. and T.C. designed the study; T.C. and X.Z. wrote the manuscript.

## Financial sources

This research was financially supported by the CAS strategic leading project ‘modular designer crop breeding’ (grant no. XDA08020301), a CAS international collaboration grant (GJHZ1501), and the National High Technology Development Program (2014AA101601).

## Competing financial interests

The authors declare no competing financial interests.

## Supplemental Data

The following materials are available in the online version of this article.

**Supplemental Data File S1.** Equations used in WACNI.

**Supplemental Figure S1.** Correlation between TGDW and 32 model parameters within the ‘natural population’ generated by WACNI.

**Supplemental Figure S2.** Prediction of TGDW variance within ‘natural population’ generated by WACNI by statistical modeling.

**Supplemental Figure S3.** The responses of major agronomic traits to variation of model parameters LPD, LONL, LSL and LINA, respectively.

**Supplemental Figure S4.** The responses of major agronomic traits to variation of model parameters LSS, LStS, LStD and LPS, respectively.

**Supplemental Figure S5.** The responses of major agronomic traits to variation of model parameters LINU, GCD, GStS and GSU, respectively.

**Supplemental Figure S6.** The responses of major agronomic traits to variation of model parameters GPS, GONU, CStS and GStD, respectively.

**Supplemental Figure S7.** The responses of major agronomic traits to variation of model parameters CPS, CPD, RINU and RINL, respectively.

**Supplemental Figure S8.** The responses of major agronomic traits to variation of model parameters RONL, RSU, RONU and RINA, respectively.

**Supplemental Figure S9.** The responses of major agronomic traits to variation of model parameters RG, RLG, RLR and XONP, respectively.

**Supplemental Figure S10.** The evolution of maximal total grain dry weight (TGDW) throughout generations in the ‘evolutionary populations’ generated by WACNI.

**Supplemental Figure S11.** Molecular and macroscopic physiological features of *in silico* high-yield rice lines under different conditions.

**Supplemental Figure S12.** Daily weather data of the normal condition used in WACNI.

**Supplemental Table S1.** Description of parameters used in WACNI.

**Supplemental Data File S2.** List of abbreviations and their definitions.

**Supplemental Data File S3.** List of major hypotheses used in WACNI.

## ACKNOWLEDGEMENTS

This research was fnancially supported by the CAS strategic leading project ‘modular designer crop breeding’ (grant no. XDA08020301), a CAS international collaboration grant (GJHZ1501), the National High Technology Development Program (2014AA101601), and the Bill and Melinda Gates Foundation Project Realizing Improved Photosynthetic Effciency (grant no. OPP1060461). The authors acknowledge International Visiting Professorship support of SPL from the Chinese Academy of Sciences.

